# AIChatBio: An Artificial Intelligence Chatbot Model for Biological Knowledge Retrieval and Biomacromolecule Design

**DOI:** 10.1101/2025.09.11.675485

**Authors:** Elisha Liu, Chen-Yu Liu

## Abstract

Conversational agents for bioinformatics data analysis and interpretation remain largely inaccessible to the broader biological research community. This gap is especially pronounced in the current Generative AI era, which demands a paradigm shift in how researchers interact with computational tools. There is a pressing need to bridge well-established biological infrastructures and databases with the capabilities of Generative AI to democratize access to bioinformatics insights.

In this study, we present an integrated framework that connects the robust bioinformatics resources of the National Center for Biotechnology Information (NCBI) with Generative AI through a novel Artificial Intelligence Chatbot Model for Biological Knowledge Retrieval and Biomacromolecule Design, AIChatBio.

This operational model positions Generative AI as an intelligent information hub. User interactions with the chatbot are enriched by real-time data retrieval from web portal of the biological databases hosted at NCBI, which are then translated into structured inquiries toward the web applications of NCBI and bioinformatics analysis tools. These inquiries are directed toward bioinformatics analysis tools to perform tasks such as sequence alignment and primer design. Additionally, the outputs generated by these tools are interpreted by the chatbot, allowing users to gain meaningful insights without requiring deep technical expertise in bioinformatics.

To demonstrate the feasibility of this approach, we developed a prototype implementation that integrates PCR primer design using Primer-BLAST [1], literature interpretation via PubMed for general topics, and the LitVar2 for SNPs associated topics [23]. This system was built using TypeScript and the ChatGPT API combining the bioinformatics web applications from NCBI, and its source code is publicly available via GitHub and the Chrome extension is available at Chrome Web Store.

Our work highlights the potential of Generative AI to transform biological data analysis workflows, making them more intuitive, accessible, and scalable for researchers across disciplines.

## INTRODUCTION

Recent advances in large language models (LLMs), such as ChatGPT, have generated considerable interest in their potential to support bioinformatics research [2–4]. These models can interpret natural language queries, retrieve relevant information, and generate step-by-step analytical guidance. As the field progresses, there is growing emphasis on developing **multimodal conversational agents** capable of understanding both natural language and biological sequences [5–7]. Such systems could enable conversational genome analysis, personalized medicine interfaces, and broader accessibility to complex biomedical tools.

One promising direction involves the integration of LLMs with genomic data to create systems that reason across modalities. A notable example is the **Chat Nucleotide Transformer**, designed to serve as a general-purpose agent for interpreting biological sequences. These models aim to infer biological function from sequence data while remaining usable by researchers without computational backgrounds. However, despite their promise, such models must be rigorously validated before being deployed in high-stakes contexts like clinical genomics or diagnostic workflows. In biomedical applications, **interpretability and trust** are essential: black-box generative AI risks misinterpreting signals or generating plausible yet incorrect hypotheses.

Several practical challenges persist. The **cost of retraining or fine-tuning LLMs** remains high, which limits accessibility for smaller research groups. Moreover, many existing approaches underutilize established biological infrastructures—such as curated databases, ontologies, and toolkits—leading to redundant or disconnected solutions.

Beyond these implementation barriers, there are significant theoretical challenges. Biological data are often **high-dimensional, noisy**, and **non-Euclidean**, while human language is **contextual, ambiguous**, and **culturally embedded**. Let biological data lie on a manifold *M*_*B*_ ⊂ *R*^*n*^and language data on another manifold *M*_*L*_ ⊂ *R*^*m*^. The challenge is to learn a mapping: *f*: *M*_*B*_ → *M*_*L*_or a joint embedding: ∅: *M*_*B*_ × *M*_*L*_ → *R*^*d*^ that preserves both semantic and structural integrity. This is nontrivial due to differences in curvature, topology, and sparsity between these data manifolds [8–11]. A joint embedding enables **cross-modal retrieval and contextual fusion** by projecting heterogeneous data into a shared vector space.

Complicating matters, biological datasets are **expensive to generate** (e.g., cryo-EM, RNA-seq) and often difficult to annotate. Annotation requires expert domain knowledge and is subject to **inter-annotator variability**. Consequently, many real-world scenarios are characterized by **low-resource learning**, which undermines generalization across modalities. In addition, many biological concepts lack direct linguistic analogs. For instance, **protein–ligand binding affinity** may not map to any single word or phrase. Domain-specific ontologies like **Gene Ontology (GO)** or **SNOMED CT** are hierarchical and semantically dense, complicating integration into LLMs. Embedding such structured knowledge typically requires **graph-based representations** [12,13], where biological entities and their relationships are modeled as a graph *G* = (*V, E*) *V*=biological entities as nodes, *E*=relations as edges. Learning embeddings: *v*_*i*_ ∈ *R*^*d*^for each node *v*_*i*_ that aligns with language embeddings *w*_*j*_ ∈ *R*^*d*^remains an **open problem in graph–text alignment** [14].

To address some of these limitations, **retrieval-augmented generation (RAG)** has emerged as a powerful strategy for enhancing LLM performance [15–17,22]. Instead of retraining or fine-tuning, RAG enables models to retrieve external information dynamically, improving **accuracy, contextual relevance**, and **factual consistency**. While multimodal RAG systems are gaining traction, most applications remain restricted to textual, visual, or audio domains [8,18–19], with limited implementation in biological formats such as sequence data or experimental metadata. Moreover, while RAG commonly relies on **vector embedding**, recent studies propose **embed-ding-free RAG** as a viable alternative [20], enabling real-time, index-free retrieval that mirrors the flexibility of human reading.

In this study, we propose **AIChatBio**, an Artificial Intelligence Chatbot Model for Biological Knowledge Retrieval and Biomacromolecule Design as a prototype demonstrating how LLMs can be applied to real-world bioinformatics workflows. By integrating existing NCBI tools and databases with LLM-based conversational interfaces, AIChatBio allows users regardless of coding or bioinformatics expertise to perform tasks such as **PCR primer design, SNP variant exploration**, and **literature search** via natural language. This model showcases the potential for Generative AI to serve as an **intelligent interface layer**, improving accessibility, reproducibility, and usability in biological research.

### The Architecture of the Operational Model

Most bioinformatics tools and pipelines require specific commands or structured keywords to perform analyses. These inputs can be effectively generated through natural language conversations between users and large language models (LLMs). With well-crafted prompts, an LLM can translate a user’s natural language query into the appropriate commands or formats required by bioinformatics applications. However, in many cases, the LLM’s output still requires further **parsing, validation**, or **formatting** before it can be directly submitted to downstream tools.

Once the LLM generates a response in the expected format, the application interface can automatically detect and process it. This triggers a pipeline where parsed commands are routed to the appropriate bioinformatics tools or web portals. These tools then perform the requested tasks— such as sequence alignment, primer design, or variant lookup, retrieve the necessary data, and return the output. The processed results are finally interpreted by the LLM and delivered to the user in a clear and actionable format.

The full workflow is illustrated in **Figure 1** and consists of the following stages:

#### 1. Inquiring

The user initiates a conversation with the LLM through a front-end application interface, expressing their query in natural language.

#### 2. Monitoring

The application continuously monitors the interaction between the user and the LLM. A pre-configured system prompt is applied to instruct the LLM to provide responses in a specific format. This format includes predefined labels or triggers that indicate when a response is ready to be processed.

#### 3. Parsing

When the LLM’s output includes the expected trigger label, the application detects it, extracts the relevant information, and parses it. The system then formulates a structured query tailored for a specific bioinformatics tool or database (e.g., NCBI’s Primer-BLAST, LitVar2, or PubMed).

#### 4. Submitting

The structured query is submitted to the target bioinformatics application, web API, or data portal for execution.

#### 5. Executing

The external tool or database processes the request, performs the necessary computation or data retrieval, and returns the result to the application. This step serves as the **retrieval** phase of the pipeline, enriching the LLM’s knowledge base with real-time information.

#### 6. Response Delivery

The application reformats the output from the bioinformatics tool into a prompt as **augmentation** phase, which is sent back to the LLM. The LLM interprets the results and generates a userfriendly, context-aware response. This stage is the **generation** phase of the pipeline, concluding the user’s inquiry cycle.

**Figure 1.**
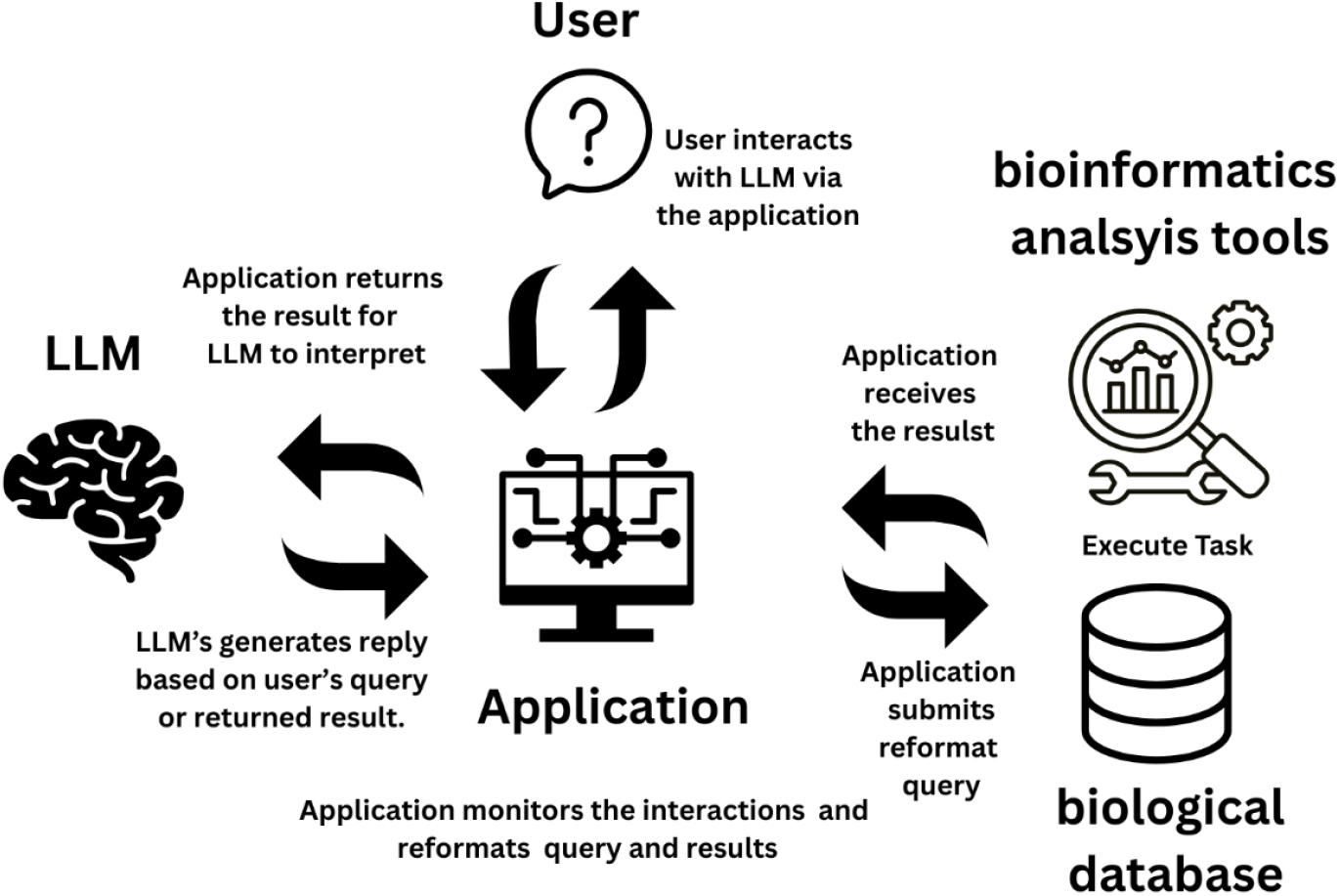
Illustration of the integrated workflow involving a user, large language model (LLM), application interface, bioinformatics tools, and biological databases.

The user initiates a query (Inquiring), which is processed by the application through prompt formatting and monitoring (Monitoring and Parsing). A structured query is then submitted to an external bioinformatics tool or database (Submitting). The tool executes the analysis and returns results (Executing), which are reformatted and sent back to the LLM via the application. The LLM interprets the results and delivers the final output to the user (Response Delivery), completing the cycle.

## IMPLEMENTATION DETAILS

To demonstrate the real-world applicability of the proposed model, we developed a Chrome Extension called **AIChatBio**.

### I. Prompt Design and Validation for AIChatBio Features

In the process of implementing the AIChatBio model, carefully engineered prompts were created to guide ChatGPT’s responses across four key functionalities:

1. **PCR Primer Designer**
2. **Primer-BLAST Navigator**
3. **Research Reporter**
4. **Genome Curator**

To support the **PCR Primer Designer** feature, one of the primary components, we applied the **Chain-of-Thought (CoT)** prompting technique [21]. This approach encourages large language models to generate intermediate reasoning steps, thereby improving accuracy and coherence when performing complex tasks.

The workflow begins when the user initiates a query via a conversational interface with ChatGPT. In response, ChatGPT returns a structured reply following a predefined format, typically including essential elements such as the **target gene** and **organism**. The AIChatBio Chrome Extension monitors the model’s output and automatically extracts the labeled information. This parsed data is then submitted to the **NCBI web portal** (e.g., Primer-BLAST) for further processing.

Once the external query is executed, the NCBI portal returns the results to AIChatBio, and the extension continues making follow-up inquiries until the task is completed. These results are then packaged with a **prompt**, which guides ChatGPT to interpret the analysis output. Finally, ChatGPT generates a coherent, human-readable summary that is delivered back to the user through the AIChatBio interface (Figure 2).

**Figure 2.**
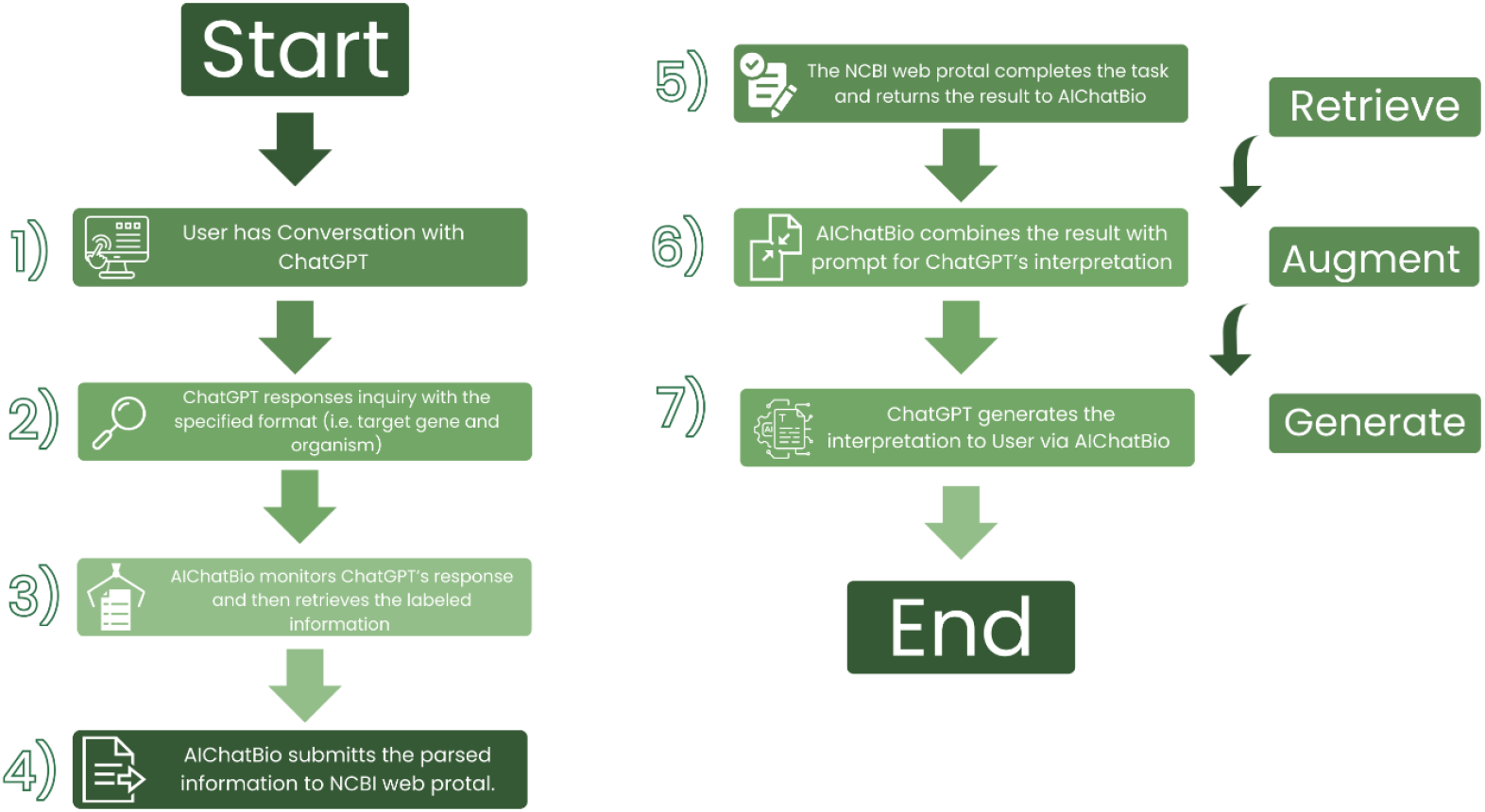
Workflow diagram illustrating the AIChatBio-assisted bioinformatics query and interpretation pipeline using ChatGPT and the NCBI web portal.

The steps are as follows: Step1. **User Inquiry**: The user initiates a request using natural language via the AIChatBio interface. Step 2. **LLM Structuring**: ChatGPT generates a structured response containing critical data such as gene name and organism. Step 3. **Monitoring and Extraction**: AIChatBio parses the labeled content from ChatGPT’s response. Step 4. **Submission to NCBI**: The parsed information is submitted to appropriate NCBI web applications (e.g., Primer-BLAST). Step 5. **Execution and Retrieval**: The web application executes the analysis and returns the results to AIChatBio. Step 6. **Contextual Augmentation**: Results are combined with a prompt that enables ChatGPT to interpret and summarize the data meaningfully. Step 7. **Final Delivery**: ChatGPT generates a user-friendly description along with a direct link for primer design, which is then presented to the user.

The right-hand side of the figure illustrates the **Retrieve–Augment–Generate** (RAG) framework employed during the interpretation process.

### II. Prompt Testing and Iterative Refinement

To refine the prompts, particularly for the **PCR Primer Designer**, we conducted iterative testing using **twenty simulated user scenarios**. Each scenario reflected different user profiles and varying levels of domain expertise—from novice researchers to advanced practitioners.

A prompt was classified as a failure if ChatGPT was **unable to identify the correct gene or organism**, returned hallucinated entities (e.g., non-existent genes), or responded with incorrect or irrelevant taxonomy terms (e.g., “Ancient Egyptian”). Other failure modes included not returning the required labeled information in the designated format, which is critical for automated parsing by AIChatBio.

This testing revealed multiple edge cases in which miscommunication or prompt ambiguity led to inaccurate or misleading outputs. Based on these findings, we iteratively revised the prompts, improving both clarity and model alignment with user intent. Below is an example of a prompt:

(*Prompt and scenario examples to follow in next section or supplementary material*.)

“The user needs to have a @@ around the identified gene and organism, but they don’t have any information on the gene and organism. In some scenarios, they have little information of the environment around the organism and the organism itself. Help them identify what gene and organism they need. Do not provide any examples of genes or organisms before they have written their scenario, that will lead them off track. In the following format, use scientific taxonomy and the gene used for the genetic test must be an encoding gene rather than other genetic marker. The exact organism or gene may not be fully identified, but use educated guesses to infer the organism and gene. Put the identified organism and encoding gene in the only one, given format (You cannot add or make any modifications to this format):

@insert identified organism@

@insert identified encoding gene@

-Also, if the user has ‘no idea”, then ask questions to find the specific organism and encoding gene. In addition, if the organism can not be identify, answer @insert unknown organism@ @insert

unknow gene@

Ground Rules:

-YOU MUST USE SCIENTIFIC TAXONOMY AND NO MITOCHONDRIAL DNA FOR THE ENCODING GENE.

-USE ONLY ONE FORMAT FOR THE ORGANISM AND ENCODING GENE

-YOU MUST USE REAL ORGANISMS AND ENCODINGS GENES. DO NOT MAKE ANY UP.”

For the second primary feature, **Primer-BLAST Navigator**, a dedicated prompt was developed to enable ChatGPT to provide real-time, contextual explanations of the Primer-BLAST interface and assist users in understanding each of its components. This feature is designed to help users— particularly those unfamiliar with the technical structure of the NCBI Primer-BLAST web application—navigate and utilize its full functionality more effectively.

To accomplish this, **HTML element IDs** from the NCBI Primer-BLAST interface were incorporated into the prompting strategy. These IDs allow the language model to recognize and associate specific interface elements with their intended functions. When a user hovers over a particular element within the interface, the system captures the corresponding HTML DOM element ID and sends it, along with a tailored prompt, to the ChatGPT API.

Upon receiving this input, ChatGPT interprets the identifier, determines the user’s area of focus, and generates an explanation of the element’s function. It also offers step-by-step usage guidance and, when applicable, suggests potential next actions. This interaction occurs in real time via the AIChatBio Chrome extension, creating a seamless, **context-aware assistance system** for bioinformatics tool navigation.

This architecture follows a **Retrieve–Augment–Generate (RAG)** framework: **Retrieve**: DOM metadata is captured based on user interaction. **Augment**: Contextual prompts are dynamically constructed. **Generate**: ChatGPT produces tailored, real-time instructions or guidance.

**Figure 3.**
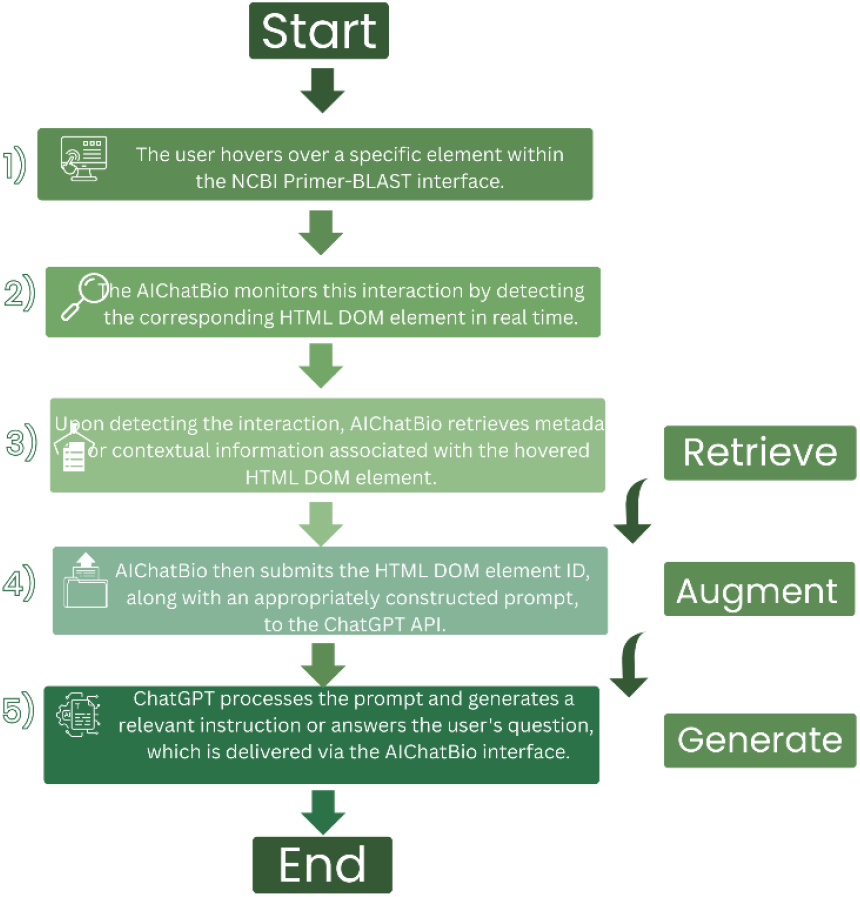
Real-time AIChatBio assistance within the NCBI Primer-BLAST interface.

**Step 1**: The user hovers over a specific element in the Primer-BLAST interface. **Step 2**: The AIChatBio extension detects the corresponding HTML DOM element. **Step 3**: Metadata or contextual cues associated with the element are retrieved. **Step 4**: This data is sent, along with a context-specific prompt, to the ChatGPT API. **Step 5**: ChatGPT processes the input and returns a context-aware explanation or instruction. **Step 6**: The output is displayed to the user via the browser extension, enabling dynamic, task-relevant assistance.

Unlike the **PCR Primer Designer**, this feature did not require extensive scenario-based testing. However, it was evaluated using **simulated user interactions**, in which test users posed questions about specific interface elements. ChatGPT’s responses were reviewed based on four main criteria:1. **Clarity and completeness** in explaining the purpose of each feature. 2.**Accuracy and relevance** of the generated answers. 3.**Topical focus**, ensuring that the output stayed aligned with the user’s intent. 4.**Practicality of suggestions**, assessing whether the guidance could lead the user toward a correct primer design outcome.

This lightweight but targeted evaluation approach helped verify that the prompt delivered consistent and high-quality performance without necessitating extensive iteration. Below is an example of a prompt designed for the **Primer-BLAST Navigator**:

“Introduction: This is the code for an NCBI primer-Blast, and you will teach the user on how to use the website to search info that I am interested.

Rules: When the user hovers over a particular element, that element will be entered into the textbox. Then, you will explain how the user can utilize that element (e.g. submit a chemical structure by clicking this button). However, that element will not be in its exact form, so it will be up to you to see which following piece of code below best matches the element that is entered into the textbox. Next, you will provide a series of suggestions of what the user should do next after inserting the information for that particular element. The users first response to you will be an element insert, so you will follow the necessary procedures as indicated in the steps. Additionally, keep your explanation simple, as possible, so that the user may understand (refrain from using complex terminology.”

Step 1. DO NOT MENTION ANY OTHER ELEMENTS ONCE THE USER CHOSE ANOTHER ELEMENT.

Step 2. Your response to this prompt should be, “What can I help you with”

Step 3. Offer suggestions of what to do next after you explain the feature to the user, or if the user doesn’t have any questions. When doing so, refer to the particular element with its ID, and not by the features name. …{Primer-BLAST page HTML elements list}..: This is the code for an NCBI Primer-BLAST, and you will teach the user on how to use the website to search info that I am interested”

**For one of the smaller features, the Research Reporter**, a dedicated prompt was designed to summarize PubMed articles retrieved from the NCBI database. AIChatBio formulates the inquiry based on the user’s keyword input and preference settings, ensuring that the request is contextually relevant and tailored to the user’s specific needs. This refined prompt is then submitted to the ChatGPT API, where it is processed to generate a concise and informative summary of the relevant literature. The response is delivered back to the user through the AIChatBio interface in a clear and readable format. *(Figure 4)*

**Figure 4.**
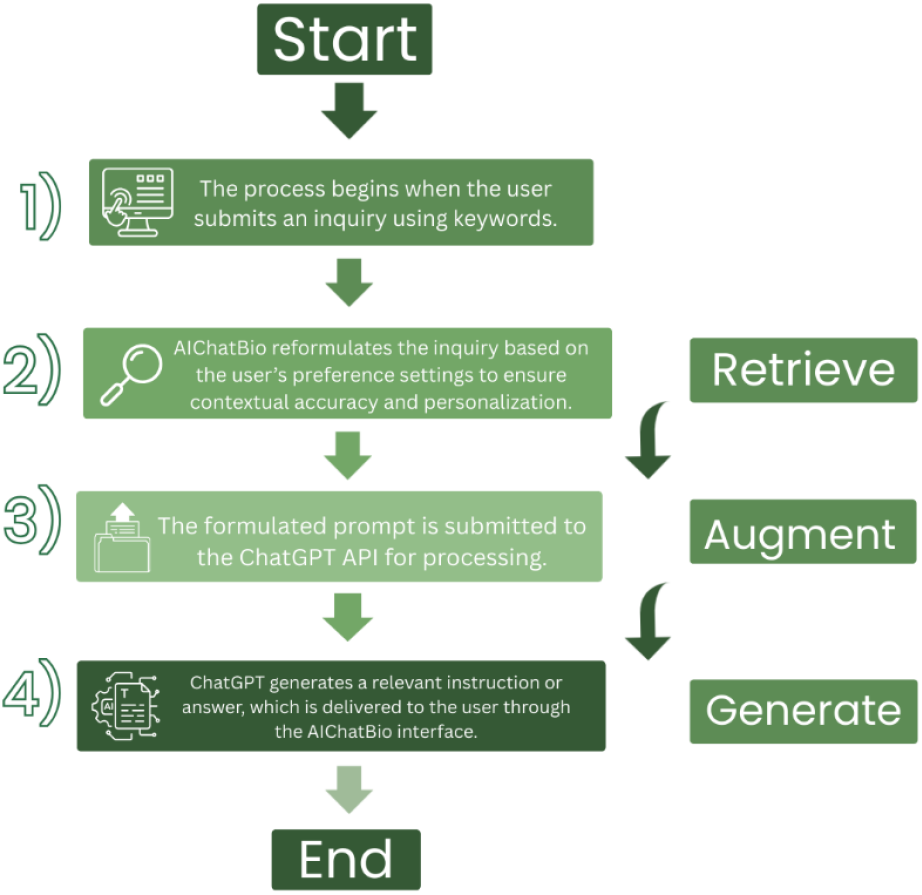
Workflow of AIChatBio-assisted keyword-based inquiry and response generation using the ChatGPT API.

Step 1: The user initiates the process by submitting a keyword-based inquiry. Step 2: AIChatBio reformulates the inquiry according to the user’s preference settings (e.g., focus area, publication recency, or summary length), ensuring contextual accuracy and personalization. Step 3: The refined prompt is submitted to the ChatGPT API for processing. Step 4: ChatGPT retrieves and synthesizes relevant information from PubMed article summaries, generating a meaningful interpretation or summary. Step 5: The AI-generated response is delivered to the user through the browser interface via the installed AIChatBio Chrome extension. Below is an example of a prompt designed for the PubMed summarization feature:

“PubMed search results: {web_results} Current date: {current_date} Instructions: Using the provided web search results, write a comprehensive reply to the given query at an {education} language level. Make sure to cite results using [<u>number</u>] notation after the reference. If the provided search results refer to multiple subjects with the same name, write separate answers for each subject. Query: {query}”.

#### Similarly, another feature—the Genome Curator—was implemented using a prompt structure similar to that of the PubMed feature

The process begins when the user submits an inquiry using any type of input related to **NCBI LitVar2**, such as a gene name, variant ID, or disease term. **AIChatBio** submits this inquiry to the **LitVar2 system**, which processes the request and returns a list of relevant **PMCIDs** (PubMed Central reference identifiers). Upon receiving the PMCIDs, AIChatBio initiates a retrieval process to access the corresponding full-text scientific articles from **PubMed Central**.

These retrieved articles are then combined with a contextually appropriate prompt, tailored to the user’s preferences (e.g., summary length, clinical relevance, or variant focus). This formulated prompt is submitted to the **ChatGPT API**, which generates a personalized and contextually accurate response, synthesizing the key information from the retrieved literature. The response is then delivered back to the user through the **AIChatBio** Chrome extension interface.

Throughout the workflow, AIChatBio ensures seamless integration of **literature retrieval** and **natural language generation**, adhering to a **Retrieve–Augment–Generate (RAG)** framework to enhance biomedical information access and interpretation. *(Figure 5)*

**Figure 5.**
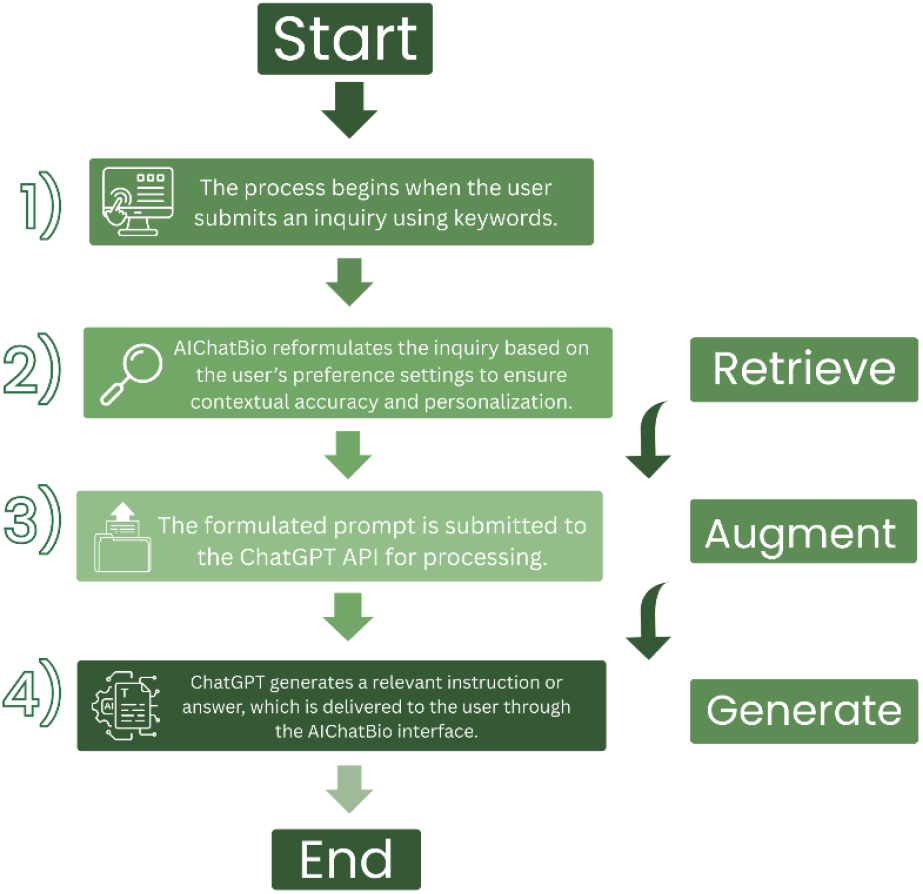
Workflow of AIChatBio-assisted literature retrieval and interpretation using NCBI LitVar2 and ChatGPT.

The process begins when the user submits an inquiry using any form of input related to NCBI LitVar2, such as a gene, variant, or disease term. AIChatBio forwards this inquiry to the LitVar2 web portal, which processes the request and returns a list of relevant PMCIDs. Based on these identifiers, AIChatBio retrieves the corresponding full-text scientific articles.

These articles are then integrated into a context-specific prompt tailored to the user’s preferences—such as desired summary depth, clinical focus, or variant relevance. This prompt is submitted to the ChatGPT API, which processes the input and generates a personalized, contextually accurate summary or explanation. The response is then delivered back to the user through the browser interface with the AIChatBio Chrome extension installed.

This workflow follows the Retrieve–Augment–Generate (RAG) paradigm, enabling efficient, real-time, and user-centered access to biomedical literature. Below is an example prompt for the Genome Curator feature:

“Var search results: {web_results} Current date: {current_date}\n\nInstructions: Using the provided web search results, write a comprehensive reply at {education_level} language level. These results are associated with the query. If possible, make a connection with the result. Make sure to cite results using [<u>number</u>] notation after the reference. If the provided search results refer to multiple subjects with the same name, write separate answers for each subject. Query:

{query}” .

### II. Technical Implementation of the ChatGPT–NCBI Extension

In addition to enabling interactions with ChatGPT, the technical architecture of the AIChatBio extension was carefully designed to ensure seamless communication between the user, the ChatGPT API, and NCBI web applications. The extension was developed using **TypeScript**, compiled into **JavaScript**, and integrated into the browser along with accompanying **JSON** and **CSS** files.

The extension is capable of capturing both the user’s input and ChatGPT’s responses directly from the **text area**, as well as from **browser elements** that the user interacts with such as through hover events. Rather than relying on the default behavior of the browser’s submit button and text area, the extension uses a **customized submit function**. This function reformats the user’s inquiry before sending it to either the **NCBI web portal** or the **ChatGPT API**, depending on the task.

In addition, the extension monitors for **label information** included in ChatGPT’s responses. When a label (e.g., @gene@) is detected, the extension parses the relevant information—such as the gene symbol and organism name—and automatically submits a query to the appropriate **NCBI web application**, such as Primer-BLAST.

This is made possible through a **DOM observer**, which continuously monitors changes in the **HTML Document Object Model (DOM)** to detect interactions and changes on the webpage. It serves as a middleware layer that facilitates communication between the **ChatGPT API** and the **NCBI web services**, ensuring the correct transfer of information and execution of tasks.

Finally, based on the user’s selection from a **dropdown menu** within the extension, the system formats the results returned by NCBI into a structure suitable for ChatGPT to interpret under its designated role (e.g., “Research Reporter” or “Genome Curator”). ChatGPT then processes these structured results and generates a final summary, which is delivered back to the user through the **AIChatBio** interface.

### II. Technical Implementation of the ChatGPT–NCBI Extension

In addition to interactions with ChatGPT, the technical implementation of the extension was carefully considered. The extension was developed in **TypeScript**, compiled into **JavaScript**, and integrated into the browser along with supporting **JSON** and **CSS** files.

The extension is designed to capture either the user’s input or ChatGPT’s reply directly from the text area, as well as from elements in the browser that the user hovers over. Instead of relying on the default behavior of the submit button and text area, the extension employs a **customized submit function**. This function reformats the user’s inquiry before posting it to the **NCBI web application portal or ChatGPT API**.

Additionally, the extension monitors any **label information** included in ChatGPT’s replies. When such a label is detected, the extension parses it and automatically submits the corresponding **gene and organism information** to the NCBI web portal. This monitoring function observes changes in the **HTML Document Object Model (DOM)** and mediates the exchange between the ChatGPT API and NCBI web applications.

Finally, depending on the settings selected in the **drop-down menu**, the customized submit function converts the results returned from the NCBI server into a format suitable for the role in ChatGPT. ChatGPT then interprets and summarizes the results based on assigned role, which are delivered back to the user.

## RESULT AND DATA ANALYSIS

To evaluate the performance of **AIChatBio**, we asked ChatGPT to randomly list 20 human genes: *BRCA1, BRCA2, TP53, EGFR, APOE, CFTR, MYC, PTEN, TNF, IL6, INS, FTO, HBB, G6PD, MTHFR, ACE2, SLC6A4, OPN1LW*, and *FOXP2*. We then compared how AIChatBio and ChatGPT handled PCR primer design for these targets. ChatGPT either declined to provide primer sequences or produced incorrect ones, as verified using the UCSC Genome Browser and UCSC In-Silico PCR (based on the GRCh38/hg38 assembly). In contrast, AIChatBio consistently returned links to validated PCR primer sets for each gene, demonstrating superior reliability and practical utility, as shown in **Figure 6**.

**Figure 6.**
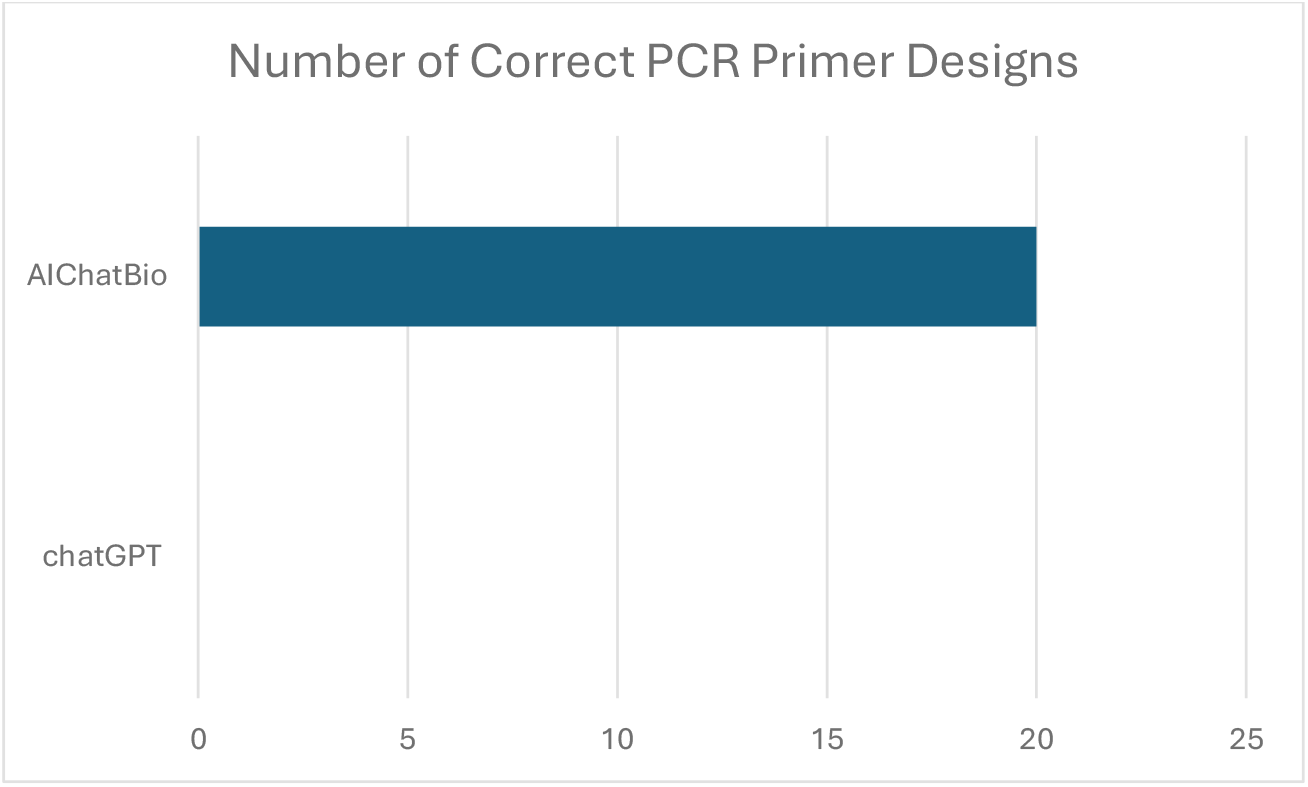
Comparison of PCR primer design performance between ChatGPT and AIChatBio. The bar chart shows the number of correct primer designs generated for 20 randomly selected human genes. ChatGPT produced 0 correct designs, either declining or generating invalid primers, whereas AIChatBio successfully returned validated primer sets for all 20 genes. This demonstrates the higher accuracy and practical applicability of AIChatBio in primer design tasks.

## DISCUSSION

Originally, our goal was to explore the feasibility of directing a large language model (LLM) through conversational interaction to perform biological data analysis. As a proof of concept for the proposed model, we developed a chatbot to assist with **PCR primer design**, since PCR (polymerase chain reaction) remains a foundational technique in molecular biology. It enables precise amplification of DNA for applications ranging from mutation detection to gene expression analysis. However, designing effective PCR primers is a nontrivial task that requires careful consideration of sequence specificity, thermodynamics, and genomic context. The process involves selecting appropriate gene targets, retrieving accurate DNA sequences, and configuring multiple design parameters—each of which can significantly influence experimental outcomes.

## RESULT and DATA ANALYSIS

### Challenges of Applying LLMs Directly to PCR Primer Design

Directly applying LLMs in this domain presents several challenges:

#### I Hallucinations and Misinformation

LLMs are prone to producing “hallucinated” outputs—confident but incorrect information. In the context of PCR, this can result in:

- Incorrect primer sequences
- Misidentified gene targets
- Invalid melting temperature or GC content calculations

#### II. Lack of Domain-Specific Constraints

Unlike dedicated primer design tools, LLMs do not inherently enforce biochemical constraints. Without structured prompts or external validation, models may:

- Ignore primer–dimer risks
- Suggest primers with poor specificity
- Overlook secondary structure interference

#### III. Task-Specific Focus Drift

LLMs are built for general-purpose conversation. Without carefully engineered prompts, they may drift away from the task, offering tangential or irrelevant information. This lack of task fidelity undermines their reliability in precision workflows like primer design.

#### IV. Limited Integration with Genomic Databases

While LLMs can describe how to use databases such as NCBI or Ensembl, they cannot directly query or validate sequence data. This limits their ability to:

- Confirm gene annotations
- Retrieve organism-specific sequences
- Cross-reference SNPs with relevant literature

Such errors may lead to failed experiments, wasted resources, and misleading conclusions.

#### V High Cost and Latency in Retrieval-Augmented Generation (RAG)

The generation phase of RAG relies heavily on LLM APIs, introducing cost and latency challenges, driven by:

- The cost of API tokens for users
- Long inference times for large models

This is particularly problematic in educational or resource-limited contexts, where frequent queries can lead to substantial operational expenses.

### Existing Tools and the Need for Conversational Interfaces

Traditional tools such as Primer3 and NCBI Primer-BLAST offer robust functionality but require technical expertise and familiarity with bioinformatics workflows. Increasingly complex designs—such as targeting single-nucleotide polymorphisms (SNPs), incorporating fluorescent probes, or working with non-model organisms—pose additional challenges for users lacking domain-specific training.

Given LLMs’ ability to translate user inquiries into meaningful input for established bioinformatics tools, we demonstrate their utility through PCR primer design by querying the NCBI database and embedding an interactive chatbot interface for NCBI Primer-BLAST.

This chatbot supports a range of functionalities, including:

- Explaining complex biochemical concepts in accessible language
- Guiding users through primer design workflows, including target selection and parameter optimization
- Suggesting relevant databases, tools, and resources based on user intent
- Assisting with literature searches related to SNPs, gene targets, and experimental protocols

### Limitations of the Current Implementation

Despite its promise, this implementation faces several limitations:

- **Token Constraints:** Due to ChatGPT’s limited context window, it cannot summarize hundreds or thousands of articles simultaneously.
- **Database Coverage:** While NCBI hosts most species and genes, some targets are absent (e.g., ancient or degraded samples like mummies).
- **Sequence Length Limits:** NCBI Primer-BLAST cannot process sequences longer than approximately 50,000 nucleotides, such as entire genomes. In such cases, the extension may return “No Sequence,” constraining user options.
- **Prompt Sensitivity:** Output quality depends heavily on prompt design. For example, requesting genome-wide primers specific to humans may exceed the model’s capabilities, leading to incomplete or misleading responses.
- **Computational Capacity:** NCBI Primer-BLAST can be resource-intensive and may struggle to handle many requests simultaneously.

These limitations highlight that the chatbot inherits not only the constraints of LLMs (e.g., token size, hallucination risk) but also those of the integrated bioinformatics resources.

## CONCLUSION

In traditional multimodal Retrieval-Augmented Generation (RAG), each modality (text, image, audio, etc.) is embedded into a vector space using dedicated models; retrieval is based on similarity within that space. The embedding-free multimodal RAG approach avoids this by using **keyword-based search and analysis commands**, leveraging structured document formats (databases, captions, tags) and rule-based principles (bioinformatics algorithms) to locate relevant content.

Applying keyword searches or analysis commands as retrievers decouples retrieval from generation, allowing context-rich documents to be passed directly to the LLM without embedding them.

The AIChatBio model demonstrates how this multimodal RAG approach can integrate biological information with natural language, and how conversational AI can bridge the gap between complex bioinformatics platforms and user-friendly interfaces. While limitations remain, iterative prompt engineering and integration with domain-specific tools enhance the reliability and accessibility of genomic analysis.

This approach holds significant potential to democratize access to molecular biology tools—particularly for students, researchers in resource-limited settings, and interdisciplinary teams—while contributing to the development of robust conversational agents for biological and medical applications.

## DATA AVAILABILITY

The full set of testing questions and scenarios as well as the prompt can be accessed through the link: https://github.com/EL-github1/aichatbio

The source code of AI ChatBio chrome extension code is also available on: https://github.com/EL-github1/aichatbio

The chrome extension is available on google web store via the link or search with the extension name AIChatBio or via the link:

https://chromewebstore.google.com/detail/oeealmonapjihcfmdkcmeakhnmphcgck?utm_source=item-share-cb

The Research Reporter in the GPT Store (with a default for summarizing three articles) is available with this link: https://chatgpt.com/g/g-VUFf5i7vY-research-reporter

The Genome Curator in the GPT Store is available with this link: https://chatgpt.com/g/g-ydaAVqWIx-genome-curator

